# Epigenetic and Transcriptomic Alterations Precede Amyloidosis in the Hippocampus of the Alzheimer’s Disease *App^NL-G-F^* Knock in Mouse Model

**DOI:** 10.1101/2025.02.20.639336

**Authors:** Mariam Okhovat, Cora Layman, Brett A. Davis, Alexandra Pederson, Abigail O’Niel, Sarah Holden, Kat Kessler, Sonia Acharya, Kandace J. Wheeler, Kimberly A. Nevonen, Jarod Herrera, Samantha Ward, Katinka Vigh-Conrad, Andrew Adey, Jacob Raber, Lucia Carbone

**Affiliations:** Department of Medicine, Knight Cardiovascular Institute, Oregon Health & Science University, Portland, OR; Department of Behavioral Neuroscience, Oregon Health & Science University, Portland, OR; Department of Molecular and Medical Genetics, Oregon Health & Science University, Portland, OR; Cancer Early Detection Advanced Research (CEDAR), Knight Cancer Institute, OHSU, Portland, OR; Division of Genetics, Oregon National Primate Research Center, Beaverton, OR; Departments of Neurology and Radiation Medicine, Division of Neuroscience, ONPRC, Oregon Health & Science University, Portland, OR

## Abstract

Detecting and understanding the early stages of Alzheimer’s disease (AD) is essential for uncovering initial mechanisms of neuropathology and devising effective interventions. In this study, we leveraged the humanized *App^NL-G-F^* mouse which exhibits early-onset amyloid pathology with a predictable timeline, to investigate molecular changes in the hippocampus and blood before the onset of severe neuropathology and independent of aging. Employing a multi-omics approach, we identified alterations in chromatin accessibility, gene expression, and DNA methylation associated with early amyloidosis. Chromatin accessibility changes were prominent in excitatory neurons during early pathology, with a later shift to inhibitory neurons, potentially reflecting compensatory mechanisms to mitigate excitatory neuron dysregulation. Despite broadly comparable hippocampal cell composition, transcriptomic comparisons between wild-type and *App^NL-G-F^* mice revealed major gene expression differences, particularly in pathways related to mitochondrial function and protein biosynthesis, preceding severe amyloid plaque deposition. In later stages, upregulation of immune and neuroinflammatory pathways was observed, aligning with established neuroinflammatory processes in AD. Additionally, we identified extensive DNA methylation differences in both the blood and hippocampus of *App^NL-G-F^*mice during early and late stages of pathology. Many differentially methylated regions in the blood, even at early pathology stages, were associated with cis-regulatory elements in the brain and were located near differentially expressed genes in the hippocampus. These regions were enriched in pathways associated with brain function, including neuron development and synaptic processes, highlighting a connection between blood methylation patterns and brain activity. This finding suggests the potential use of blood DNA methylation as a biomarker for the early detection of amyloidosis. Notably, we identified five candidate biomarker genes, including *Rbfox1* and *Camta1*, with epigenetic dysregulation detectable in both the brain and blood prior to severe amyloid accumulation. Our study, leveraging a unique AD mouse model and a multi-omics approach, highlights epigenetic signatures of AD before the onset of clinical symptoms, providing a foundation for future research into early diagnosis and therapeutic strategies, as well as potential blood biomarkers.

## INTRODUCTION

Alzheimer’s disease (AD) is a complex progressive neurodegenerative disorder and the most common cause of dementia, affecting millions of people worldwide (Knopman et al. 2021). AD is characterized by hallmark pathological features, including extracellular amyloid-β (Aβ) plaques, intracellular tau neurofibrillary tangles, neuroinflammation, and widespread synaptic and neuronal loss (Vitek 1989). Despite decades of intensive research, the precise mechanisms driving the onset and progression of AD remain elusive, including whether intracellular or extracellular Aβ causally contributes to the disease beyond being a hallmark (Small 2024), and effective therapeutic options are limited. One of the greatest challenges in AD research lies in uncovering the molecular and cellular changes that arise during the preclinical phase—a prolonged asymptomatic stage during which pathological features, such as Aβ deposition, develop (Gale 2024). This phase, which can span several decades, represents a critical window for therapeutic intervention aimed at preventing or delaying the onset of clinical symptoms. However, studying preclinical AD, presents several challenges, particularly when working with human subjects, including difficulties in identifying at-risk individuals, limited access to relevant tissues, the presence of confounding factors (*e.g.*, aging, lifestyle, and environment), variability in disease progression, and the subtlety of molecular changes at early stages of pathology. Overcoming these barriers is critical for understanding the early mechanisms driving AD and enabling the development of effective strategies to combat later symptoms.

To address these challenges, a range of genetically modified animal models, particularly transgenic mice, have been developed (Bryan et al. 2009). While no single model fully replicates the complexity of AD, each successfully mimics key features of the condition making their detailed characterization crucial for understanding AD mechanisms. In this study, we took advantage of the *App^NL-G-F^* mouse model (Saito et al. 2014) which carries a knock-in of the human amyloid precursor protein (APP) with three specific mutations: the Swedish mutation (KM670/671NL), which increases the overall production of Aβ_40_ and Aβ_42_; the Arctic mutation (E693G), which promotes the formation of insoluble Aβ fibrils; and the Beyreuther/Iberian (I716F) mutation, which raises the ratio of Aβ_42_ to Aβ_40_. The *App^NL-G-F^* model exhibits early-onset amyloid pathology starting at 2 months (∼8 weeks) of age, with neuropathology progressing until saturation by 7 months (∼28 weeks). Cognitive impairments typically manifest by 6 months (∼24 weeks) (Kundu et al. 2021; Holden et al. 2022). This predictable disease timeline makes the *App^NL-G-F^*model useful for studying molecular changes across all stages of pathology, particularly the preclinical stages. Importantly, due to the early onset of disease, the *App^NL-G-F^* model allows for the exploration of early molecular events without the confounding effects of aging, offering a robust platform for investigating early molecular events and for identifying blood-based biomarkers of brain dysregulation before the onset of clinical symptoms.

In this study, we leveraged a combination of single-cell and bulk epigenetic and transcriptomic data from a cohort of *App^NL-G-F^*mice and their wild-type littermates to investigate early molecular changes in the hippocampus, a brain region critical for memory, learning, and spatial navigation (Maguire et al. 1999) which is known to exhibit early and pronounced vulnerability in AD, with significant tissue loss and connectivity disruptions (Samuel et al. 1994). Our findings reveal substantial early epigenetic and gene expression dysregulation in the hippocampus of *App^NL-G-F^* mice prior to the onset of severe amyloidosis. Importantly, we identified epigenetic signals in blood that could potentially serve as biomarkers for early detection of neuropathology, paving the way for blood-based diagnostic tools in preclinical AD.

## RESULTS

### Hippocampal cell composition remains largely stable during progression of amyloid pathology

To investigate changes preceding and following onset of severe amyloidosis in the brain and blood of the *App^NL-G-F^* mouse model, we collected cortex, hippocampus, and blood from 30 *App^NL-G-F^* and 30 wild-type littermates. The cohort included equal numbers of male and female, sampled at postnatal week 3 (weaning), 8, and 24 (n=5 per sex/genotype/age combination; **Fig. 1A**). All assays conducted in this study (**Fig. 1B**) were performed on this cohort of animals. We selected postnatal week 3 (W3) to represent an early pre-symptomatic stage, prior to the onset of severe Aβ pathology to allow for the investigation of initial molecular and cellular changes preceding severe disease.

**Figure 1.**
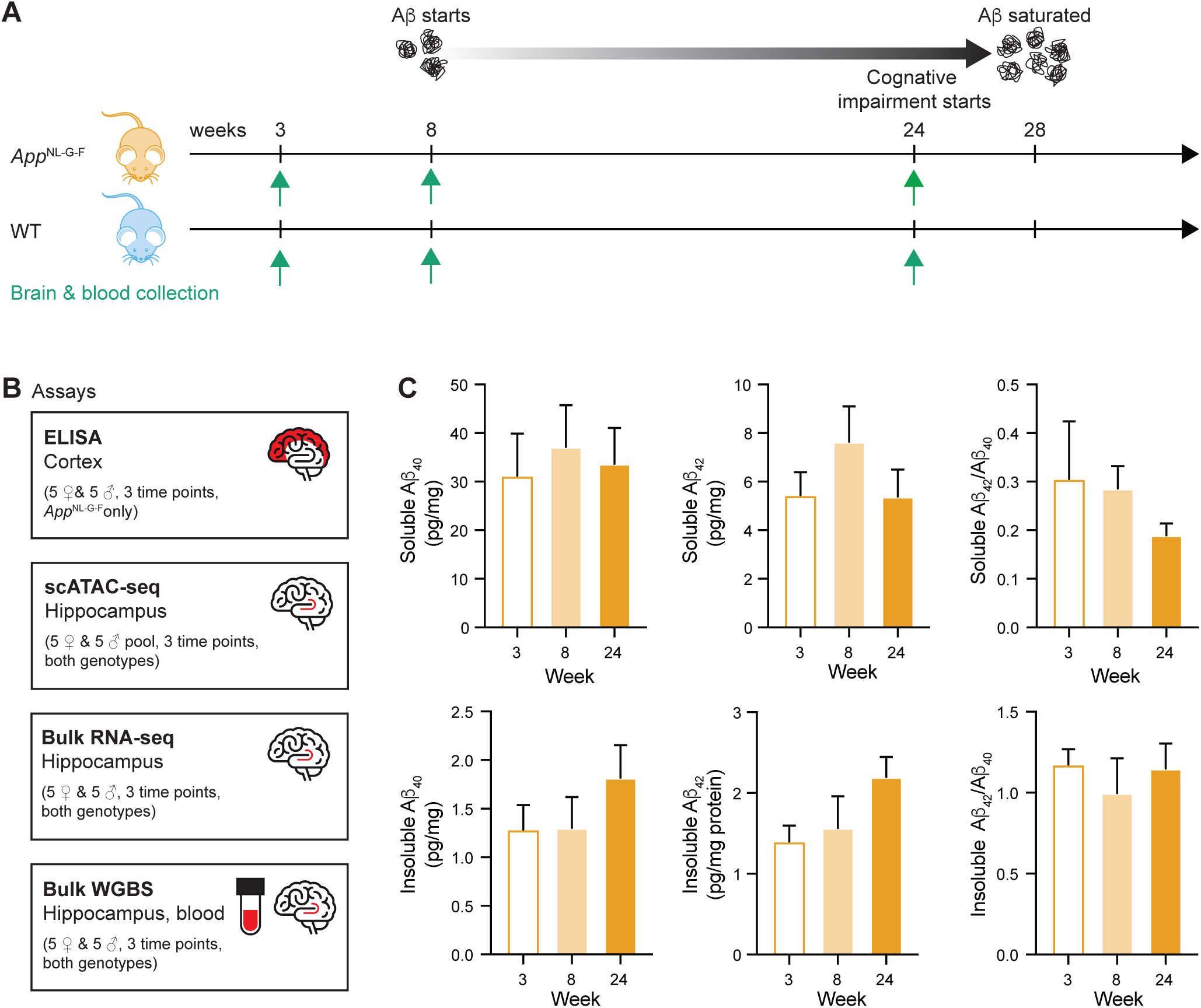
The App^NL-G-F^ mouse model allows studying Aβ pathology along a predictable timeline. **A)** The timeline of Aβ pathology in the App^NL-G-F^ model and timepoints investigated in this study. **B)** A summary of all assays conducted in this study. **C)** Bar graphs show mean soluble an insoluble cortical Aβ_40_ and Aβ_42_ levels as measured by ELISA. Error bars indicate SEM.

After performing ELISA on cortex of *App^NL-G-F^* to track the levels of soluble and insoluble Aβ_40_ and Aβ_42_ across the three time points (**Fig. 1C)**, no sex differences were seen in soluble and insoluble cortical Aβ_40_ and Aβ_42_ levels, motivating us to combine the sexes in the downstream analysis (n=10 per age). At W3, *App^NL-G-F^* mice already exhibited detectible levels of soluble and insoluble Aβ_40_ and Aβ_42_ in the cortex, signaling an even earlier onset of amyloidosis than previously reported (Saito et al. 2014). By postnatal week 8 (W8), while levels of insoluble Aβ_40_ and Aβ_42_ remained similar to W3, levels of soluble Aβ_40_ and Aβ_42_ increased, reflecting ongoing disease progression (**Fig. 1C**). At week 24 (W24), levels of insoluble Aβ_40_ and Aβ_42_ rose, indicating increased Aβ deposition, while soluble Aβ_40_ and Aβ_42_ remain relatively unchanged compared to W3. Of note, the ratio of soluble cortical Aβ_42/40_ decreases progressively with age, suggesting changes in Aβ production or clearance dynamics as the disease advances. In contrast, the insoluble cortical Aβ_42/40_ ratio remained relatively stable across ages, pointing to consistent deposition patterns of both Aβ forms over time (**Fig. 1C**).

Changes in brain cell composition, including neuron loss and gliosis, have been observed in AD (Johnson et al. 2021). To investigate changes in brain cell-composition and chromatin accessibility in the *App^NL-G-F^* mice during progression of Aβ pathology, we used hippocampus single-cell ATAC-seq (scATAC-seq) to study *App^NL-G-F^* (n=5 per sex) and wild-type littermates (n=5 per sex) at W3, W8 and W24. Given the lack of reported sex differences in amyloidosis patterns in the *App^NL-G-F^* model (Saito et al. 2014) and our own findings showing similar cortical Aβ_40_ and Aβ_42_ levels across sexes, data from males and females were combined to improve statistical robustness in all analyses. To improve cell yields, nuclei from biological replicates were pooled to create one pool per sex/genotype/age combination yielding a total of 12 pools, comprising 14,907 cells. To identify and characterize cell types within our scATAC-seq dataset, we conducted unsupervised clustering and identified 21 cell clusters (**Fig. 2A**). Cross-reference of cluster-specific accessible chromatin regions with known marker genes (**Fig. 2B**; **Supplementary** Fig. 1, (Johnson et al. 2021; O’Connell et al. 2023) identified the following cell-types: microglia (n=615 cells), oligodendrocyte (n=1,729), precursor/immature oligodendrocyte (n=349), endothelial (n=127), astrocyte (n=771), Cajal-Retzius cells (n=63), excitatory neurons (n=8,070) and inhibitory neurons (n=1,362) (**Fig. 2B; Supplementary Table 1**). Although most clusters were successfully annotated, four (C8, C9, C10 and C19) could not be confidently classified due to ambiguous and inconsistent chromatin accessibility patterns (**Fig. 2B**; **Methods**). These clusters, which comprise a total of 471 cells, likely represent rare or transitional cell types (C8 and C10), as well as technical noise (C9 and C19) and were therefore excluded from downstream analysis.

**Figure 2.**
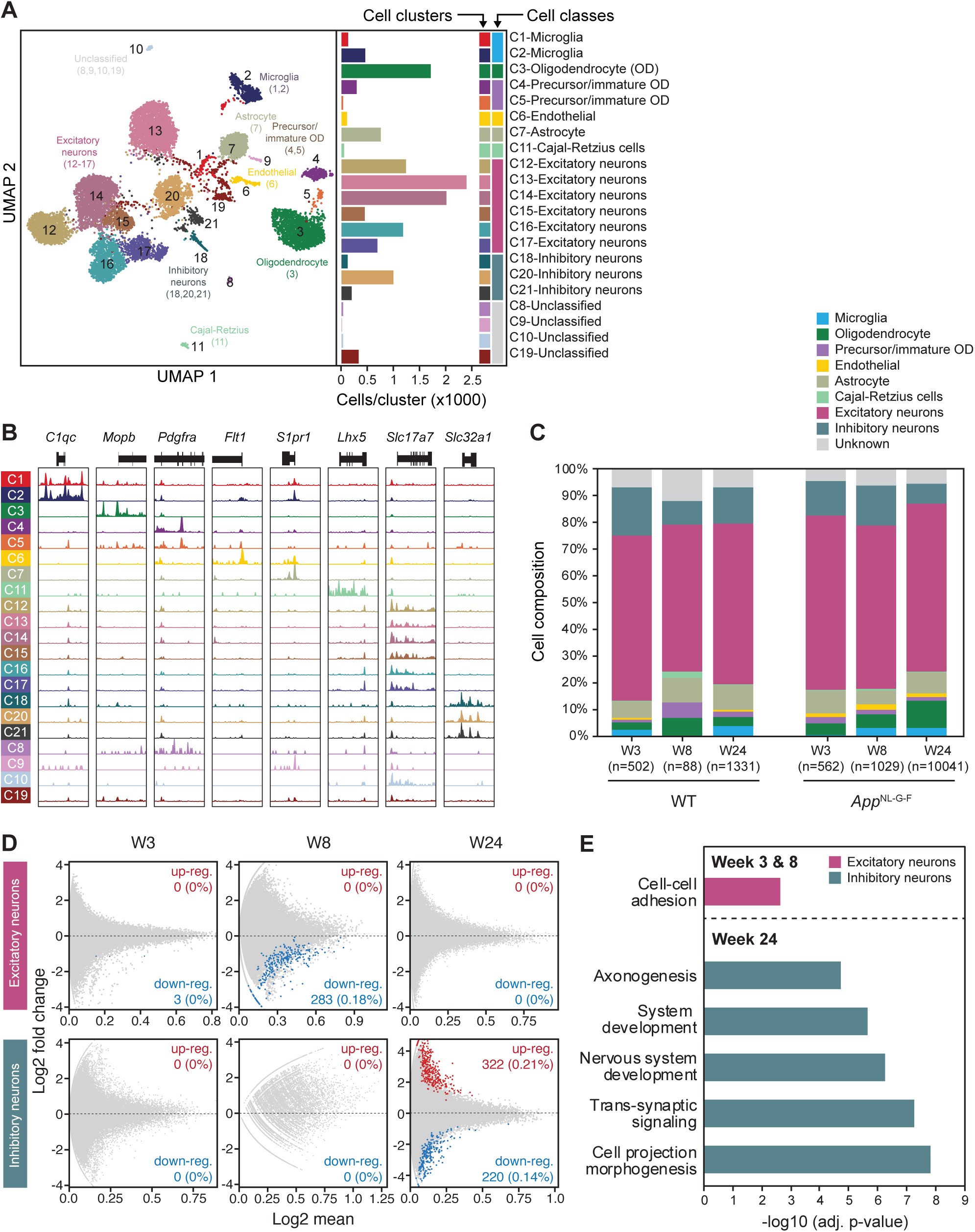
scATAC-seq reveal cell-specific changes in chromatin accessibility in early vs. late Aβ pathology. **A)** Left: uniform manifold approximation and projection (UMAP) dimensionality reduction after iterative LSI of scATAC–seq data from 12 pooled samples. Each dot represents a single cell (n = 14,907), colored by its corresponding cluster. Right: Bar plot shows the number of cells per cluster. Bar plot showing the number of cells per cluster, with corresponding cluster colors and assigned cell type **B)** Genomic tracks display chromatin accessibility at a subset of marker genes used to annotate cell types in this study. assignments. **C)** Estimated hippocampus average cell composition is shown for each time point and genotype. **D)** MA plots of differential scATAC-seq peaks reveal that chromatin accessibility differences are apparent in excitatory neurons during early pathology (W3 and W8), while in late pathology (W24), these differences are exclusive to inhibitory neurons. **E)** Significant gene ontology (GO) terms associated with differentially accessible regions in corresponding cell types during disease progression.

We used cell distribution across clusters to estimate cellular composition of the hippocampus across ages and genotypes (**Fig. 2C**; **Supplementary Table 1**). Across all samples, excitatory neurons were the most prevalent cell type (61.0%± 0.44; mean ± SEM), followed by inhibitory neurons in most cases (12.6% ± 0.73%; mean ± SEM). While drastic shifts in cell composition were not observed between genotypes and across time points, we detected a mild increase in microglia and oligodendrocyte abundance in *App^NL-G-F^* mice with time (**Fig. 2C**), which is consistent with the increased amyloidosis (**Fig 1C**) and likely reflects heightened neuroinflammation. We also noted a slight decrease in the proportion of neurons with age progression in both *App^NL-G-F^* and wild-type animals. In WT animals, average neuron composition declined from 79.8% to 73% between W3 and W24, and in *App^NL-G-F^* it decreased from 78.1% to 70.1%, indicating a modest loss in neuron proportion as mice transition from weaning to adulthood. Additionally, in W8 wild-type mice, we noted deviations in composition of some cell types (*e.g.*, microglia and precursor/immature oligodendrocytes; (**Fig. 2C**). However, these are likely due to the low number of cells recovered from this pool of samples (n=88), compared to the other time points (2,693 ± 4121; mean ± STDEV; **Supplementary Table 1**), and given the consistent cell composition observed at W3 and W24 (**Fig. 2C**), it is reasonable to assume W8 would have shown a similar profile with higher cell counts. In general, we observed only subtle shifts in cell composition between genotypes and time points, and such shifts seem to be in line with the physiological changes occurring with AD progression in our model.

### Chromatin accessibility changes first appear in excitatory neurons, but shift to inhibitory neurons later in amyloidosis

To identify condition-specific patterns in chromatin accessibility, we first performed marker peak analysis and identified scATAC-seq peaks unique to each age/genotype/cell-type condition (q<0.1). At W3, significant marker peaks were identified exclusively in excitatory neurons (n=6), while at W24, most marker peaks were found in inhibitory neurons (n=37; **Supplementary Table 1**). The scarcity of marker peaks in other cell types is likely due to the low and inconsistent number of cells representing those cell types in our dataset (**Supplementary Table 1**). Marker peaks corresponding to W3 excitatory neurons in *App^NL-G-F^* mice overlapped with several genes relevant to AD pathology and neurodegeneration, such as the mammalian target of rapamycin (*mTOR*) and *Smpd3* (Stoffel et al. 2018; Baloni et al. 2022).

Considering that marker peaks were predominantly detected in excitatory and inhibitory neurons, we performed follow-up differential scATAC-seq analysis only for these cell types (**Supplementary Table 1**). Consistent with our previous findings, in early stages of pathology (W3 and W8), we only detected significantly different *App^NL-G-F^* vs WT chromatin accessibility peaks in excitatory neurons (n=3 at W3, and n=283 at W8; FDR<0.1). All these regions exhibited reduced accessibility in *App^NL-G-F^*mice compared to WT (**Fig. 2D**). Gene ontology (GO) analysis of W3 and W8 differential peaks in excitatory neurons revealed significant enrichment of biological pathways relevant to cell-cell adhesion (q<0.05; **Fig. 2E; Supplementary Table 2**), possibly indicating early impairment in connectivity and signaling of excitatory neurons. Furthermore, several of the differential peaks overlapped genes implicated in AD or neurodevelopment. For example, one of the earliest differential peaks in excitatory neurons was found at the promoter of the *Kdm6a* gene, which codes for an X-linked H3K27 histone demethylase whose dysregulation disrupts neurodevelopment (**Supplementary** Fig. 2**; Supplementary Table 1**; (Lindgren et al. 2013; Van Laarhoven et al. 2015)). Among other AD-relevant genes at W8, we detected reduced accessibility at the promoter of *Picalm* (log2fold change= −2.07, FDR=0.06) and at putative intronic enhancers in *Itm2b* (log2fold change= −1.28, FDR=0.08) and *Ngfr* (log2fold change= −3.51, FDR=0.08) genes (**Supplementary** Fig. 2). *Picalm* (Phosphatidylinositol binding clathrin-assembly protein), which in humans is the most significant genetic AD susceptibility locus after *APOE* and *BIN1*, is involved not only in Aβ production and clearance, but also in tau-mediated neurodegeneration and in endocytic homeostasis (Ando et al. 2022). *Itm2b* (Integral membrane protein 2B gene), which is implicated in the regulation of excitatory synaptic transmission at both pre- and post-synaptic termini, has been implicated in familial AD (Yao et al. 2019), and *Ngfr* (Nerve growth factor receptor) induces neurogenic plasticity and is a major player in AD pathology (Siddiqui et al. 2023). These findings suggest that early chromatin accessibility changes at key AD-related genes in excitatory neurons are associated with, and may contribute to, the onset of amyloid neuropathology.

In contrast to W3 and W8, differential accessibility peaks at W24 were completely absent from excitatory neurons and only present in inhibitory neurons (n= 542; **Fig. 2D**; **Supplementary Table 1**) and were almost equally split between regions of increased (∼69%) and decreased accessibility (∼41%) in *App^NL-G-F^*. GO term analysis revealed enrichment of several pathways related to neuron projection, neurogenesis, and synaptic signaling (**Fig. 2E; Supplementary Table 2**), indicating major changes in function of inhibitory neurons during later amyloidosis. Several of the differential accessibility regions at W24 overlapped directly with AD-relevant genes (**Supplementary Table 1**). For example, inhibitory neurons of *App^NL-G-F^* mice exhibited higher accessibility at a putative enhancer located in the intron of *Optn* (Optineurin) gene (log2fold change= 4.34, FDR=0.07; **Supplementary** Fig. 2), a receptor that increases expression of autophagic genes to reduce neurotoxicity and inflammation in AD (Xu et al. 2022). Also, *App^NL-G-F^* mice showed reduced accessibility at a putative intronic enhancer at the *Ptk2b* gene, also known as *Pyk2* or Protein tyrosine kinase 2 beta (log2fold change= −2.48, FDR=0.02; **Supplementary** Fig. 2), which is a susceptibility gene for late-onset Alzheimer’s disease. PTK2B is implicated in AD through its involvement in synaptic plasticity, calcium signaling, and neuroinflammatory pathways, all of which are disrupted in AD (Heneka et al. 2015; Skaper et al. 2017; Chami 2021). Across the three disease stages we only found one gene, *Epb41l1* (Walensky et al. 1999), that showed differential accessibility at both early (W8) and late (W24) pathology in excitatory and inhibitory neurons, respectively (**Supplementary** Fig. 2; **Supplementary Table 1**). However, the regions do not perfectly overlap, and the direction of change is opposite, with *App^NL-G-F^* excitatory neurons showing reduced accessibility at W8 while inhibitory neurons displayed increased accessibility at W24.

The *EPB41L1* gene, which encodes the Band 4.1-like protein 1, links cell adhesion molecules to G-protein coupled receptors at the neuronal plasma membrane and has been associated with neurofibrillary tangles in AD brains (Sihag et al. 1994). All in all, our scATAC-seq results highlight a shift in epigenetic dysregulation from excitatory to inhibitory neurons as AD pathology progresses, with differential accessibility peaks increasingly associated with genes involved in neurogenesis, synaptic signaling, and neuroinflammation.

### Significant gene expression changes in the hippocampus of *App^NL-G-F^* mice precede neuropathology

The overall stability we observed in hippocampal cell composition across genotypes and ages alleviates concerns that differential gene expression in bulk tissue could be drastically skewed by cell composition differences that have been previously observed in AD (Johnson et al. 2021; Yap et al. 2024). Thus, to investigate gene expression changes in the hippocampus before and during AD pathology, we obtained RNA-seq data from bulk hippocampus tissue of the same *App^NL-G-F^* and wild-type mice (**Fig. 1A, B**). We found that the number of differentially expressed genes (DEGs) between *App^NL-G-F^* and WT mice varied drastically across the three timepoints. The highest number of DEGs were identified at W3 (n=2,628; adjusted p≤0.05), despite low insoluble cortical Aβ_42_ levels at this age, suggesting complex gene regulatory changes occur even before amyloid accumulation and neuropathology. DEGs at W3 were evenly split between being over-expressed (49.7%; n=1,306) and under-expressed (50.3%; n=1,322) in *App^NL-G-F^* mice (**Fig. 3A**; **Supplementary Table 3**). The numbers of differentially expressed genes (DEGs) were much lower at W8 (n=4) and W24 (n=156), even though insoluble cortical Aβ_42_ levels at W8 were comparable to W3 and higher at W24 (**Fig. 1C**). The four DEGs identified at W8 were all under-expressed in *App^NL-G-F^*hippocampus compared to controls, while at W24 nearly all DEGs (94%; n=147) were over-expressed (**Fig. 3A)**. Across disease stages, only 17 DEGs were shared between W3 and W24 (**Fig. 3A**), with eight of these exhibiting opposite expression changes between the two time points, suggesting they are differentially affected in early versus late amyloid pathology. Altogether, the large number of DEGs identified at W3, along with the variable gene expression patterns detected across ages, indicate that major gene dysregulation precedes severe neuropathology, and that the brain undergoes distinct physiological states as AD progresses.

**Figure 3.**
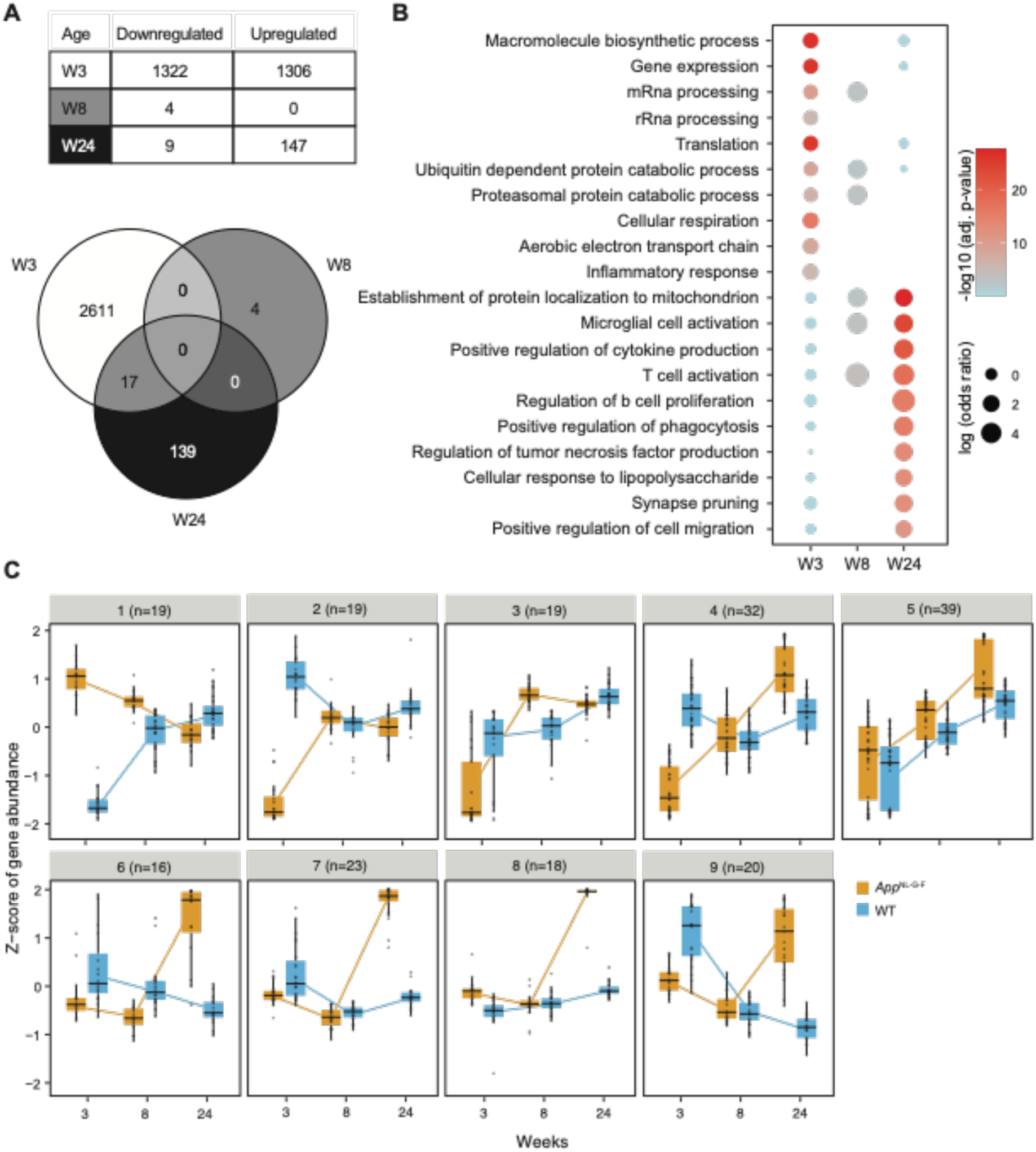
Gene expression changes in App^NL-G-F^ hippocampus precede severe Aβ pathology and highlight brain dynamics during disease progression. **A)** A table summarizes differentially expressed genes (DEGs) across timepoints of pathology and a Venn diagram illustrates DEGs shared across timepoints **B)** Selected significantly enriched gene ontology (GO) terms for DEGs at each timepoint highlight key pathway shifts during disease progression. **C)** Clustering of genes with significant genotype-by-age interactions reveals major trajectories of gene expression changes throughout disease progression.

### Gene expression profiles reveal distinct pathways involved in pre- and post-AD pathology

To shed light on the physiological states of the *App^NL-G-F^*brain as it transitions from exhibiting no symptoms to severe amyloid pathology, we set to identify over-represented biological pathways among differentially expressed genes (DEGs; q≤0.05) at each time point using EnrichR (Chen et al. 2013) (**Supplementary Table 4**). DEGs identified at W3, which comprised of both upregulated and downregulated genes, were enriched in several biological pathways, even though insoluble cortical Aβ_42_ levels were lowest at this stage. Significantly enriched biological pathways (q<0.01) included major pathways of protein biosynthesis and catabolism including “gene expression”, “mRNA splicing”, “translation”, and “proteasomal protein catabolic process”, likely reflecting alteration in gene expression paradigms before or in response to the earliest traces of amyloidosis. The rest of significant pathways were mostly relevant to “mitochondria function and organization” (**Fig. 3B**; **Supplementary Table 4)**, consistent with previous reports suggesting mitochondria and metabolic dysregulation as one of the earliest mechanisms leading to AD pathology (Moreira 2010; Wang et al. 2020). Many of the DEGs corresponding to these pathways have been previously implicated in cellular respiration (*e.g.,* components of the electron transport chain, such as *Ndufa8 and Cyc1*), or mitochondrial organization (*e.g.,* translocase of inner mitochondrial membrane genes, such as *Timm23* and *Timm44*), indicative of extensive disruption of mitochondrial function in our model preceding amyloid neuropathology. At W8, the number of DEGs (n=4) was too small to provide reliable ontology enrichment. However, DEGs at W24, which were mostly upregulated in *App^NL-G-F^*, revealed several significant biological pathways, predominantly associated with immune and neuroinflammatory response (**Fig. 3B; Supplementary Table 4**). Notable genes included those implicated in microglia activation (*e.g., Trem2*), phagocytosis (e.g. *Tlr2*) and cytokine signaling (e.g. *Csf1r*). Additionally, DEGs at W24 were significantly associated with synapse pruning, aligning with the cognitive decline observed in the *App^NL-G-F^* model around this age (Paula I. Moreira 2010; Wang et al. 2020).

We also identified 209 genes with significant “age-by-genotype interaction” (q<0.05), representing genes whose expression is affected differently during aging in *App^NL-G-F^* versus WT mice, and clustered them based on their expression trajectories (**Fig. 3C**; **Supplementary Table 5**). Most of these genes were also identified as a DEG at least one age point (**Supplementary Table 4**), showing a dynamic relationship with age in *App^NL-G-F^*mice. Specifically, 121 of the genes with significant interaction were differentially expressed at W3, one at W8, 72 at W24, and 12 at both W3 and W24. Among the gene clusters with significant age-by-genotype interaction, two clusters (clusters 1 and 2) contained genes with highly divergent *App^NL-G-F^*expression at W3, exhibiting opposite age-dependent changes in expression between genotypes (**Fig. 3C; Supplementary Table 5**). Given the early and distinct expression divergence of these genes, they may be involved in the onset and progression of neuropathy. While we did not detect significant enrichment of biological pathways among the genes in these clusters (q<0.05), they did include several genes implicated in neurodegeneration, such as *Crebbp*, *Atxn2*, and *Mt3*(Rouaux 2004; Koh and Lee 2020; Laffita-Mesa et al. 2021). Four other clusters (clusters 6, 7, 8, and 9) represented genes whose expression trajectory culminated in highly divergent expression at W24 (**Fig. 3C; Supplementary Table 5**). These genes were enriched in several immune response pathways, including B-cell proliferation, dendritic cell antigen processing/presentation, and phagocytosis (**Supplementary Table 5**). This suggests that the immune response follows a markedly different trajectory and reaches a highly divergent state in brains predisposed to APP pathology.

### DNA methylation patterns in brain and blood of *App^NL-G-F^* mice differ from wild-types and are linked to hippocampal gene regulation

We used whole genome bisulfite sequencing (WGBS) to investigate epigenetic alteration in the hippocampus and blood of *App^NL-G-F^* mice throughout disease progression, and to determine if methylation changes in the blood are associated with gene dysregulation in the brain. To explore the role of DNA methylation in disease progression of the *App^NL-G-F^* model, we first identified significant differentially methylated regions or DMRs (difference in methylation ≥10% and q ≤0.05) in hippocampus (HPC-DMRs) and blood of *App^NL-G-F^* mice relative to WT (**Supplementary Table 6**). To better infer the functional roles of these DMRs, each was assigned to the nearest gene’s transcription start site, assuming it to be the most likely regulatory target. In the hippocampus, we identified 537 significant DMRs between *App^NL-G-F^* and wild-type mice at W3, 276 at W8 and 510 at W24. In the blood, there were 664 DMRs at W3, 3,476 DMRs at W8 and 262 DMRs at W24 (**Fig. 4A; Supplementary Table 6**). Approximately 5% of genes associated with hippocampal DMRs in the brain, and 4% of DMR-associated genes in the blood were shared across two or all three time points in the same tissue, suggesting some persistence of DNA methylation dysregulation within each tissue throughout disease progression (**Fig. 4A**). In the hippocampus, over half of the DMRs were hypermethylated in each time point (**Fig. 4B**). The overlap of hippocampal DMRs with genomic features showed consistent patterns across all three time points, with a nearly equal distribution between genic and intergenic regions. Among the genic DMRs, most overlapped intronic regions, while the majority of intergenic DMRs were non-coding, suggesting they overlap gene regulatory elements (**Fig. 4C**). In the blood, DMRs at W3 and W24 showed broadly similar methylation changes and genomic overlaps as DMRs in the hippocampus. However, blood DMRs at W8 were more numerous, mostly hypomethylated and predominantly genic, with nearly half overlapping exons (**Fig. 4A and 4C**).

**Figure 4.**
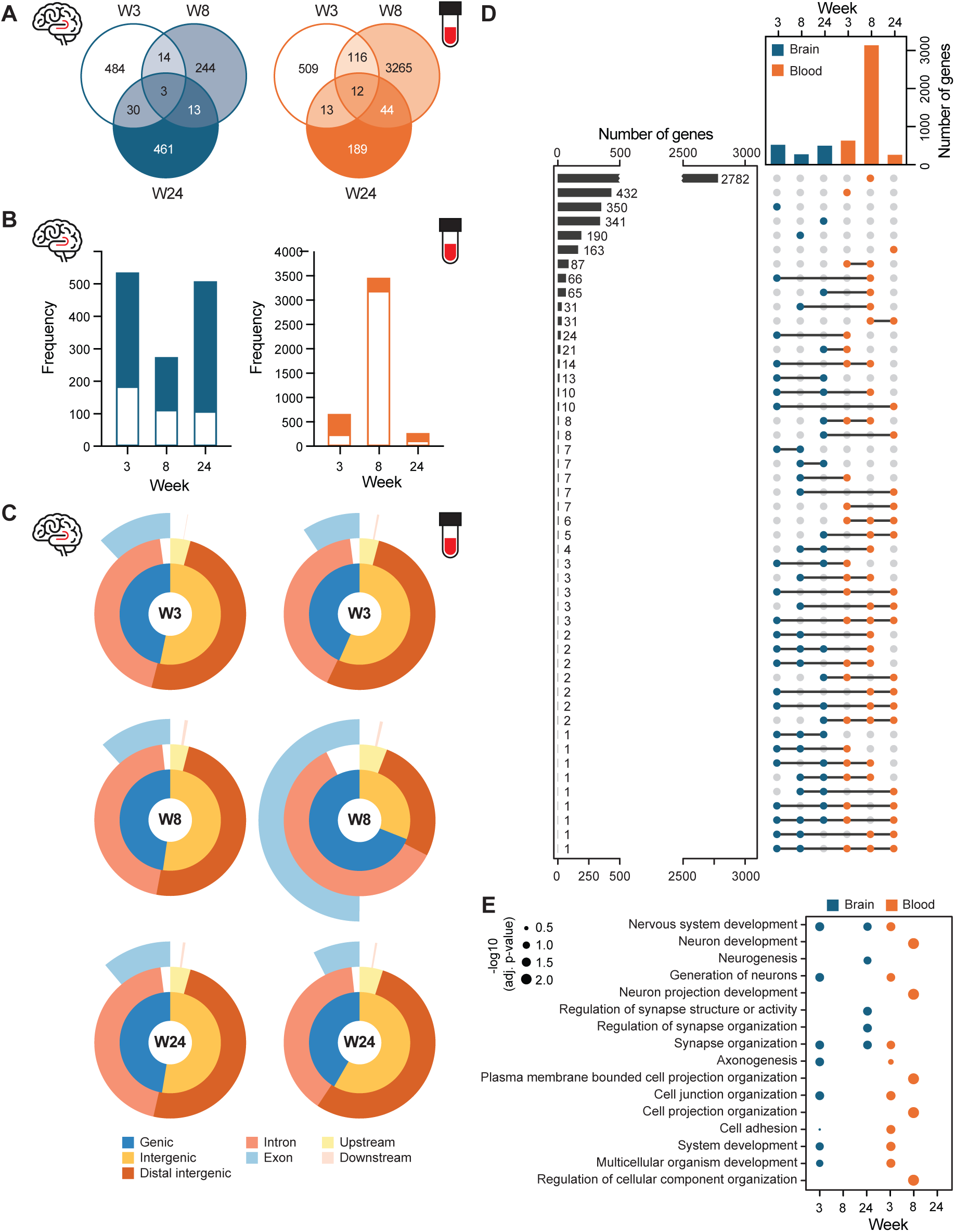
App^NL-G-F^ mice show dynamic DNA methylation changes in hippocampus and blood. **A)** Venn diagram shows DMR-containing genes shared across different timepoints in blood and hippocampus. **B)** Bar graphs show proportion of hypermethylated and hypermethylated DMRs across all ages in hippocampus and blood. **C)** Pie charts show disruption of DMR overlapping various gene features. Intergenic DMRs located >3kb from the start or end of a gene were classified as “distal intergenic”. Intergenic DMRs located <3kb from a gene start or end were categorized as being “upstream” or “downstream”, respectively. **D)** Upset plot shows number of gene associated with at least one DMR across three timepoints and two tissues, illustrating the extent of overlap between these groups. **E)** Dot plots display the statistical significance of the top five most enriched biological Gene Ontology (GO) pathways associated with DMRs in the hippocampus and blood.

We next evaluated the correspondence between DNA methylation in the hippocampus and the blood. Overall, we found average gene promoter methylation in blood and brain of individuals to be significantly correlated regardless of age and genotype (**Supplementary** Fig. 3). We identified several genes that were differentially methylated in both hippocampus and blood (although their DMRs were not necessarily located at the same genomic locations). Most of these genes showed differential methylation in the blood at W8 (**Fig. 4D**), likely due to the unusually high number of blood DMRs at this time. Notably, W3 showed the second-highest number of DMR-associated genes shared between the brain and blood (**Fig. 4D**), indicating the potential to detect early blood-based epigenetic signals that reflect brain states during the initial stages of disease progression. When we restricted the analysis to DMRs overlapping the exact same genomic regions, rather than those associated with the same genes, we identified only seven shared DMRs, with two detected at the same age (**Supplementary Table 6; Supplementary** Fig. 4). One of these, is a 1 kb non-coding DMR located approximately 15 kb upstream of the *Tmprss15* gene, displaying 32% hypomethylation in the brain and 37% hypomethylation in the blood of *App^NL-G-F^* mice at W3. This hypomethylation remains significant in the blood by W8. (**Supplementary Table 6; Supplementary** Fig. 4). *Tmprss15* is thought to play a role in neurogenesis and/or APP metabolism and has been found duplicated in some early-onset Alzheimer’s disease cases associated with Down syndrome (Wiseman et al. 2015). Despite containing DMRs, *Tmprss15* does not show differential expression in our bulk RNA-seq data. Another DMR shared between blood and brain was observed at W8 overlapping the *Eif4A3* gene (**Supplementary** Fig. 4). This region is 11% hypomethylated in blood, but 11% hypermethylated in brain. Eif4A3 is an RNA-binding protein implicated in RNA metabolism, splicing, and nonsense-mediated decay (Ye et al. 2021), as well as axon development (Alsina et al. 2024), processes which may be linked to broader mechanisms of neurodegeneration. The rest of the DMRs shared between brain and blood were associated with *Mettl27*, *Tns1*, *Srp54b*, *Gm32357* and *Rai2* genes (**Supplementary Table 6; Supplementary** Fig. 4), which to our knowledge, lack strong ties to neurodevelopment or neurodegenerative disease.

Lastly, we investigated the potential relationship between DNA methylation and gene regulation in the hippocampus. Across all three time points and in both blood and hippocampus, we observed a significant negative correlation between promoter DNA methylation levels and hippocampal gene expression (**Supplementary** Fig. 5). This pattern suggests that DNA methylation in both the blood and brain is broadly associated with gene regulation in the hippocampus, regardless of genotype. Based on a published single-cell atlas of adult mouse cerebrum *cis*-regulatory elements (i.e. CREs) (Li et al. 2021), we found that 42% of hippocampus W3 DMRs, 44.2% of W8 DMRs and 38.2% of W24 DMRs overlap with at least one CRE identified in one or more mouse brain cell types. Similarly, in the blood, 33.6% of DMRs at W3, 75.5% of DMRs at W8, and 28.2% at W24 overlapped with brain CREs (Li et al. 2021), indicating that methylation changes detected in blood and hippocampus of *App^NL-G-F^*may impact function of brain gene regulatory elements. Additionally, the distance between differentially expressed genes in the hippocampus of *App^NL-G-F^* mice and the nearest DMRs in either blood or brain was significantly shorter than expected by chance (Wilcoxon signed-rank test, p <0.05). All in all, these observations indicate that DNA methylation differences in *App^NL-G-F^*brain and blood are tied to genotype-dependent alterations in hippocampal gene regulation and expression. Gene ontology analysis of hippocampal DMRs revealed notable enrichment in pathways related to neurodevelopment, synapse organization and cell projection at both early (W3) and late (W24) disease stages (Benjamini-Hochberg adjusted p <0.05; **Fig. 4E**). In line with the limited number of DEGs at W8, hippocampus of *App^NL-G-F^* mice also exhibited the fewest DMRs at W8, with no significant gene ontology pathways identified at this stage. Of note, blood DMRs at early stages of neuropathology (W3 and W8) also showed significant enrichment in pathways related to neurogenesis and cell projection (Benjamini-Hochberg adjusted p <0.05), but no significant pathways were recovered for blood DMRs at W24 (**Supplementary Table 7; Fig 4E**). Altogether, these observations suggest that epigenetic dysregulation associated with neuronal function persists through stages of neuropathology and that early epigenetic alterations in the blood could reflect neuronal dysfunction in *App^NL-G-F^* mice, highlighting the potential utility of blood methylation as biomarkers for detecting early amyloidosis.

### Integration of multi-omics data identifies potential early blood-based epigenetic biomarkers for APP pathology

To identify candidate biomarkers for early detection of APP pathology in the blood, we focused on the W3 timepoint to capture signals in the earliest stages of neuropathy—before onset of major histological or cognitive symptoms. We established stringent criteria to maximize reliability and generalizability of our candidate biomarkers: First, the candidate gene must contain a DMR in the blood of *App^NL-G-F^* mice at W3, allowing for early epigenetic detection in an accessible tissue. Second, the candidate genes should contain at least one DMR in the hippocampus at W3, though exact sequence overlap between blood and brain DMRs is not necessary. Third, the candidate gene linked to DMRs should be differentially expressed in the brain of *App^NL-G-F^*mice at W3.

Through the integration of our datasets, we identified five genes —*Rbfox1*, *Camta1*, *Diaph2*, *Mkl2* and *Manea—* that exhibit significant epigenetic changes in the blood and corresponding dysregulation in the brain at W3, making them strong candidates for early blood-based biomarkers (**Fig. 5A**). Of note, two of these genes—*Rbfox1, Camta1*—are also implicated in neurodevelopment and/or neuropathology, with *Rbfox1* particularly standing out due to its documented role in early and preclinical AD (Raghavan et al. 2020). In our study, blood of *App^NL-G-F^* mice showed a 1 kb region of significant hypomethylation in an intron of *Rbfox1*, overlapping a putative mouse brain CRE (**Fig. 5B**; (Li et al. 2021)). Average methylation at this DMR did not correlate with expression of *Rbfox1* in the hippocampus across individuals (**Fig. 5B**). However, *App^NL-G-F^* mice showed significant, though mild, heightened expression of *Rbfox1* in the hippocampus at W3 (log2 fold change = 0.15, FDR = 0.05; **Supplementary Table 3**) along with an intronic hypomethylated DMR, distinct from the blood DMR (**Fig. 5B**). *Camta1*, a calcium-responsive transcriptional regulator highly expressed in the brain, showed hypomethylated intronic DMRs in both hippocampus and blood at W3 (**Fig. 5C**), with mild expression increase in the hippocampus (log2 fold change = 0.15, FDR = 0.02; **Supplementary Table 3**; (Bas-Orth et al. 2016)). Of note, average methylation at the *Camta1* W3 blood DMR was significantly correlated with hippocampal *Camta1* expression across all individuals (*r*= 0.35, p=0.04; **Fig. 5C**). While no clear link exists between *Diaph2* and AD, methylation at the W3 *Diaph2* blood DMR was positively associated with hippocampal expression across all individuals (r=0.42, p=0.027; **Fig 5D**), indicating a functional link between this DMR and hippocampal gene expression. All five genes identified as potential biomarkers display additional hyper-and/or hypomethylated DMRs at W8 and/or W24, hinting at prolonged and cross-tissue dysregulation of these genes during amyloidosis progression.

**Figure 5.**
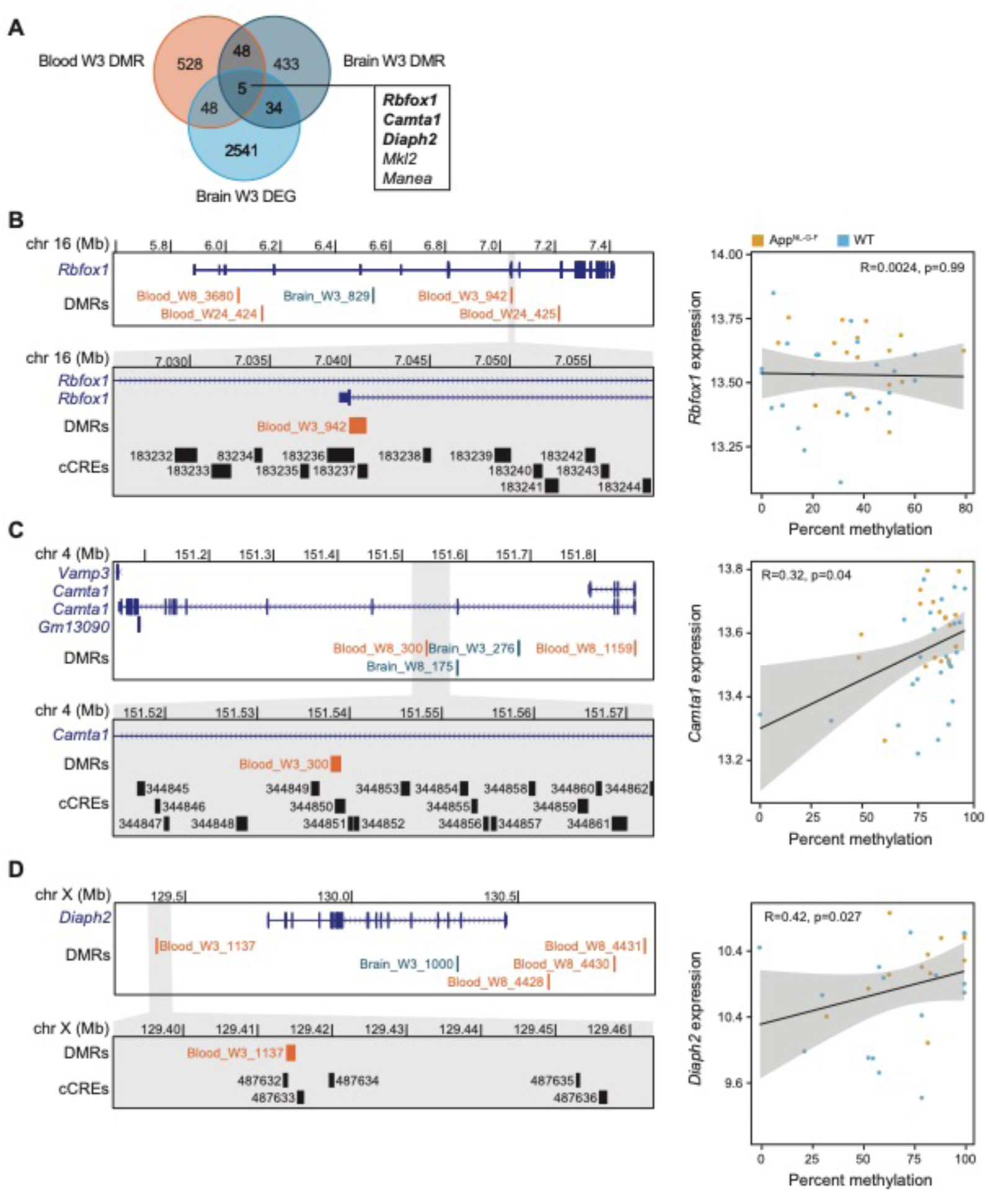
Identification of candidate early blood-based biomarkers through integration of gene expression and DNA methylation data. **A)** Venn diagram displays the number of genes meeting the criteria for candidate early blood-based biomarkers for amyloidosis. **B)** Left: UCSC genome browser screenshots provide a zoomed-out and zoomed-in view of the candidate biomarker DMR associated with Rbfox1, alongside additional brain or blood DMRs identified at other time points. The cCREs track displays mouse brain cis-regulatory elements from Li et al. 2021. Right: A scatterplot shows the average methylation at the focal DMR versus the normalized expression of the corresponding gene in the hippocampus. **C-D)** Similar UCSC genome browser views and methylation versus expression plots are shown for the candidate biomarkers associated with Camta1 (C) and Diaph2 (D).

It should be noted that our data integration also identified many additional candidate genes that met only a subset of our stringent criteria for biomarkers. Several of these genes were highly relevant to AD and/or neurodevelopment (**Supplementary Table 8**). For instance, among the 48 genes containing DMRs in both the blood and brain of *App^NL-G-F^* mice at W3 but were not DEGs, at least six have been widely implicated in neurodevelopment and neurological diseases: *Abca13, Adgrl3, Cntnap2, Dab1, Dpp6* and *Lingo2* (Knight et al. 2009; Vilarino-Guell et al. 2010; Poot 2015; Maussion et al. 2017; Nawa et al. 2020; Moreno-Alcazar et al. 2021). Additionally, of the 48 candidate marker genes differentially methylated in the blood and differentially expressed in the brain at W3, several are linked to neurodevelopment and neuropathology, including *Anks1b, Atxn2, Auts2, Dcc, Gabbr2, Hdac, Nfia,* and *Slc16a2* (Oksenberg and Ahituv 2013; Yamamoto et al. 2014; Yoo et al. 2017; Carbonell et al. 2019; Li et al. 2020; Laffita-Mesa et al. 2021; Dini et al. 2023; Singh et al. 2024). Overall, our findings highlight five strong blood biomarker candidates, as well as many additional genes of interest that merit further investigation as potential early biomarkers or contributors to AD pathology.

## DISCUSSION

Understanding Alzheimer’s disease (AD) during its preclinical stages is essential for uncovering the early mechanisms driving neuropathology and identifying biomarkers for early diagnosis and intervention. However, studying AD at these stages presents significant challenges, including difficulty in accessing relevant tissues (e.g., brain) during preclinical stages and the inability to predict who will develop AD and when. These obstacles make capturing the disease early inherently complex, necessitating innovative approaches and animal models. In this study, we take advantage of the *App^NL-G-F^* mouse model, which develops early-onset amyloid pathology along a predictable timeline without overexpression of APP beyond physiological levels and without the presence of murine APP (**Fig. 1A**). This model provides a controlled system to investigate the mechanisms preceding severe AD neuropathology, independent of the confounding effects of advanced aging. We used both male and female samples as replicates to enhance statistical power. While this approach may overlook potential epigenetic sex-specific differences, it does not compromise our primary aim of identifying universal markers of early pathology applicable to both sexes.

Despite the progressively worsening amyloid deposition reported previously (Saito et al. 2014; Masuda et al. 2016; Holden et al. 2022), and confirmed here, we found hippocampal cell composition of the *App^NL-G-F^* mice to remain relatively stable across W3, W8 and W24 timepoints, and to broadly resemble that of wild-type (WT) counterparts. The small changes that were detected with disease progression, such as neuronal loss and increased glial cells, align with reports from human AD brains (Malpetti and al. 2020), but were less pronounced in our study, likely due to the earlier pathological stages we investigated, the relative age difference of our mice and AD patients, and/or species differences in susceptibility to develop AD pathology. This cellular stability allowed us to investigate changes in gene expression and DNA methylation in bulk tissue, without major confounding effects from cell type shifts. It should be noted however, that the absence of major changes in cell composition does not rule out functional alterations within cell populations, as supported by numerous changes we detected in chromatin accessibility, gene expression, and DNA methylation.

Despite the overall stability in cell composition, the hippocampus of *App^NL-G-F^* mice showed changes in chromatin accessibility with age, specifically in neuronal cells. Other cell types, particularly glial cells, such as microglia and astrocytes, likely also undergo significant changes in our model during disease progression, as observed in human AD studies (Smith et al. 2022). However, their detection may have been hindered by the low and variable non-neuronal cell counts in our scATAC-seq assay. In early pathology stages (W3 and W8), we noted most chromatin accessibility changes in excitatory neurons, suggesting excitatory neurons to be more susceptible to early Aβ deposition, aligning with previous findings in a tau pathology mouse model (Fu et al. 2019), reports of selective neuronal vulnerability in neurodegenerative diseases (Morrison et al. 1998), as well as a higher hippocampal activity and pathological neuronal hyperexcitability in earlier stages of AD (Putcha et al. 2011; Kamondi et al. 2024). These early epigenetic changes in excitatory neurons may reflect the initial stages of cellular stress, or compensatory attempts to maintain synaptic function and plasticity despite increasing Aβ burden. Later in pathology (W24), an epigenetic shift was evident, with inhibitory neurons showing the most substantial changes in chromatin accessibility. These changes may reflect the increasing cellular stress due to Aβ burden and worsening neural network dysfunction, which likely contributes to the cognitive impairment observed in the *App^NL-G-F^* model around this age (Saito et al. 2014). Epigenetic modifications in inhibitory neurons may represent compensatory mechanisms aimed at maintaining inhibitory control and mitigating excitatory neuron dysregulation and potential excitotoxicity (Palop et al. 2006; Harris et al. 2020). Overall, the transition of impact from excitatory to inhibitory neuronal changes during disease progression underscores the dynamic interplay between Aβ pathology and the disruption of the delicate balance between excitatory and inhibitory neuronal activity, a key contributor to cognitive deficits in AD (Palop et al. 2006; Harris et al. 2020). These findings also signify the importance of understanding temporal and cell-type-specific mechanisms in AD for developing effective and targeted therapeutic strategies. Based on our observations, targeting epigenetic and gene expression changes in excitatory neurons during early AD stages may offer therapeutic benefits by preserving their function and preventing downstream effects on neural circuits. In later stages, therapies aimed at supporting inhibitory neuron function and mitigating their dysregulation could help stabilize neural networks and slow disease progression.

Surprisingly, the most extensive gene expression differences we observed in the hippocampus of *App^NL-G-F^* mice were at W3, despite the lowest cortical amyloid deposition at this age. This finding suggests that major molecular dysregulation precedes the onset of severe neuropathological symptoms, underscoring the importance and promise of studying early stages of disease. The pathways enriched among these early DEGs, particularly those related to mitochondrial function and protein biosynthesis, indicate that metabolic and protein homeostasis disruptions are among the earliest transcriptional changes in the hippocampus. This is consistent with several previous reports on AD patients (Olagunju et al. 2023). The lack of significant DEGs at W8 may be attributable to technical limitations in our study. However, if validated by further research, this finding could indicate that W8 represents a transitional stage in disease progression, during which compensatory mechanisms begin to fail, giving way to neuroinflammation and synaptic dysfunction. Further studies focused on this timepoint could help clarify the molecular events happening during this period. Consistently, at W24, when amyloid deposition is the most pronounced, the changes in expression profile mostly showed enrichment of immune and neuroinflammatory pathways. The upregulation of genes associated with microglial activation, phagocytosis, and cytokine signaling at this later stage aligns with the high level of Aβ plaques reported at this age and the well-documented role of neuroinflammation in later stages of AD pathology (McGeer and McGeer 2010). The observed changes in synapse pruning pathways at W24 further align with the cognitive deficits reported in the *App^NL-G-F^*model at this age, suggesting that the immune response may play a role in synaptic loss and dysfunction during AD progression. Overall, the variations in gene expression patterns in early versus late amyloidosis suggest that the brain undergoes drastically different physiological phases as symptoms emerge and worsen.

Our whole-genome bisulfite sequencing revealed that significant DNA methylation differences also exist between *App^NL-G-F^* and wild-type mice, spanning all stages of disease in both the hippocampus and blood. In the hippocampus, differentially methylated regions in early (W3) and late (W24) stages showed enrichment of neurodevelopment and neuron connectivity pathways. These findings suggest persistent epigenetic dysregulation linked to hippocampal neuronal function throughout disease progression. Interestingly, hippocampus of W8 *App^NL-G-F^* mice exhibited the fewest DMRs and no significant pathway enrichment, supporting the idea that this stage may represent a transitional phase, a hypothesis warranting further investigation. In the blood we observed dynamic methylation changes across timepoints, with a striking increase in DMRs at W8. Although direct overlaps between blood and brain DMRs were rare, the broad associations between blood methylation and hippocampal gene regulation, overlap of blood DMRs with brain cis-regulatory elements, and the proximity of DMRs to hippocampal DEGs, suggests that blood methylation patterns reflect gene dysregulation in the brain. Furthermore, both blood and brain DMRs were enriched in pathways related to neuronal development in the earliest stage of amyloidosis, underscoring a link between peripheral epigenetic signals in the blood and early brain dysregulation. These findings highlight the potential utility of blood DNA methylation as a biomarker for detecting early stages of amyloidosis and related neuropathological processes.

By integrating our multi-omics data, we identified DMRs at *Rbfox1*, *Camta1*, *Diaph2*, *Mkl2* and *Manea* as the most promising biomarkers exhibiting epigenetic dysregulation in both brain and blood before onset of severe amyloidosis. Notably, the current literature implicates *Rbfox1*, *Camta1* in neurodevelopment and/or neuropathology (Raghavan et al. 2020), with *Rbfox1* (RNA Binding Fox-1 Homolog 1) being particularly significant. *Rbfox1* encodes a neuronal RNA-binding protein, and recent human studies have identified it as a novel locus associated with AD during preclinical stages (Raghavan et al. 2020). In AD patients, RBFOX1 is notably localized around amyloid-β plaques and reduced RBFOX1 expression correlates with a higher amyloid-β burden (Raghavan et al. 2020) . *Camta1* (calmodulin-binding transcription activator 1) is a transcription factor regulating gene expression in response to calcium signaling. Dysregulated calcium signaling is associated with several hallmarks of AD, including amyloid-β deposition, tau hyperphosphorylation, synaptic dysfunction, and apoptosis, suggesting that CAMTA1 may contribute to AD progression (Ge et al. 2022) . Additionally, CAMTA1 is thought to regulate neuroprotective genes under stress, and significant differences in *CAMTA1* methylation have been found in the peripheral blood of stroke patients (Liu et al. 2022). Mutations or disruptions in *CAMTA1* have also been implicated in various neurological and neurodegenerative diseases, including intellectual disability, ataxia, and amyotrophic lateral sclerosis (Long et al. 2014; Wijnen et al. 2020; Alves et al. 2022). The multifunctional role of CAMTA1 in neurodevelopment and neuroprotection highlights its relevance as a candidate for further investigation in AD and amyloidosis. Moving forward, validating these candidates in human cohorts and elucidating their functional roles in early disease progression will be critical. Nevertheless, our findings underscore the potential of DNA methylation at these loci as early indicators of amyloidosis and provide a foundation for further research into the mechanisms driving their epigenetic dysregulation.

Despite the intriguing findings of our study, several limitations warrant consideration. First, the *App^NL-G-F^* mouse model used in this study mirrors familial autosomal dominant AD, which accounts for only 1-5% of all AD cases, raising questions about the generalizability of our findings to sporadic and late-onset AD. Moreover, this model does not develop tau pathology, a hallmark of AD, potentially overlooking critical amyloid-tau interactions. Furthermore, our analysis was limited to three timepoints and one brain region, which may miss key transitions in gene expression and epigenetic regulation occurring at other intervals or in other tissues. Additionally, the brain’s heterogeneous cellular composition and the differential vulnerability of cell types to AD pathology(Saura 2023) highlight the need for single cell approaches with improved resolution and sample sizes, as well as reduced technical variability. Such methods would provide more precise insights into cell-type-specific dynamics, addressing the limitations of bulk tissue analysis. Finally, given the well-documented sex differences in AD risk (Farrer et al. 1997), future studies should explore whether distinct molecular and epigenetic mechanisms underlie or precede neuropathology in males and females.

Nevertheless, our study demonstrates the power of a multi-omics approach in identifying molecular changes that precede severe neuropathology in an amyloidosis mouse model. Notably, blood-based DNA methylation changes at neurodevelopmental genes, such as *Rbfox1* and *Camta1* emerge as promising early biomarkers of amyloidosis, even before clinical symptoms appear. These findings highlight the potential for non-invasive diagnostic tools leveraging blood epigenetic biomarkers, offering a transformative step toward early detection and intervention strategies for Alzheimer’s disease. Future research should validate these findings in human cohorts and elucidate the functional consequences of these epigenetic and gene expression changes in both brain and peripheral tissues, paving the way for improved understanding and treatment of this devastating disease.

## METHODS

### Mouse breeding paradigm and tissue collection

The mice were maintained on a 12/12 h light/dark schedule (lights on at 06:00). Laboratory chow (PicoLab Rodent diet 20, #5053; PMI Nutrition International, St. Louis, MO, USA) and water were provided ad libitum. All procedures complied with the National Institutes of Health Guide for the Care and Use of Laboratory Animals and with IACUC approval at Oregon Health & Sciences University. We bred hAPP knock-in (KI) mice containing the Swedish, Iberian and Arctic mutation (i.e. *App^NL-G-F^*) on a C57BL/6J background, generated by Dr. Saito (Saito et al. 2014; Saito et al. 2016) and shared with us with C57BL/6J wild-type (WT) mice from Jax. Heterozygous breeding pairs were set up to generate homozygous *App^NL-G-F^* and WT littermates. We genotyped all offspring and only used homozygous *App^NL-G-F^* and WT individuals for downstream analysis. The genotyping protocols are available on the Riken Institute web site (Saito et al. 2014; Saito et al. 2016). Briefly, we used primers reported in Table 1 to amplify part of exon 17 containing the Artic mutation. The PCR products were digested with the MboII restriction enzyme and genotype was determined based on digestion fragments (WT genotype is cut into 171 bp and 67 bp fragments, while presence of the Arctic mutation prevents digestion). For genotyping of the Iberian mutations, we used PCR to amplify part of exon 17 containing the Iberian mutation (**Table 1**). The PCR products were digested with the BsaBI restriction enzyme (WT amplicon is cut into 171 bp and 67 bp fragments, while presence of the Iberian mutation prevents digestion). Due to COVID19-related modified operations, we shipped some biopsy samples for genotyping to Transnetyx, Cordova, TN.

**Table 1.**
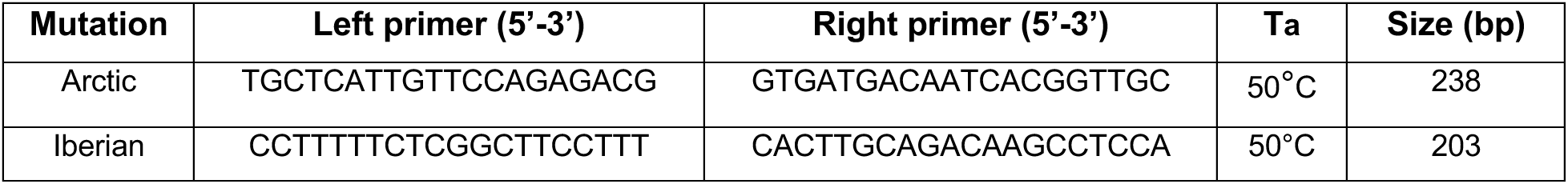
Primers used for genotyping.

At postnatal week 3, 8, or 24 (W3, W8 or W24), we collected five females and five males per genotype (homozygous *App^NL-G-F^* vs. WT), for a total of 60 individuals. The mice were euthanized by cervical dislocation. Trunk blood was collected into EDTA-treated tubes. Blood was centrifuged at 5,500 g for 10 min and the supernatant was transferred to a new tube and stored at −80°C until assay. The hippocampus from both hemispheres was dissected. Each hippocampus sample was briefly homogenized and split into three portions before freezing and storage at −80 °C.

### Cortical Aβ40 and Ab42 Enzyme-Linked Immunosorbent Assay (ELISA)

Fresh frozen cortices of the same 60 subject described above were processed for analyses of soluble and insoluble Aβ40 and Aβ42 levels using MyBiosource ELISA kits (catalog # MBS760432 and MBS268504, respectively; Thermo Fisher Scientific) according to the recommended guidelines. To each thawed cortical tissue sample comprised of both hemispheres, we added 400μl of buffer A, which is phosphate-buffered saline containing a protease inhibitor tablet (cOmplete™, 11836170001 Roche, Millipore Sigma). Next, the tissue was homogenized using a Polytron for 10 sec, followed by sonication and centrifugation at 45,000 rpm for 20 min at 4°C. The supernatant was collected as the soluble fraction. The same volume of buffer A was used to loosen the pellet. The sample was centrifuged again at 45,000 rpm for 5 min at 4°C. After removing the supernatant in a separate tube. the pellet was dissolved in Buffer B (containing 6 M Guanidine H-Cl and 50 mM Tris) and incubated at room temperature for 1 h. After this incubation, the sample was sonicated for 20 sec and the extracted pellet was centrifuged at 45,000 rpm for 20 min at 4°C. The supernatant was collected as the insoluble fraction. Pilot experiments were performed to determine the optimal sample dilution. Standard curves were generated with the same buffer dilution as the samples. The ELISAs were read at 450 nm using a SpectraMax iD5 Multi-Mode Microplate Reader (Molecular Devices). The standard curves were generated and the levels in the samples determine using GraphPad Prism software v10.2. Total protein amounts in the samples were determined by a BCA protein assay kit (Pierce, Thermo Scientific, catalog #23225) and by reading the samples at 562 nm using the iD5 Reader.

### Single-cell assay for transposase-accessible chromatin sequencing (scATAC-seq) library preparation and analysis

To obtain sufficient nuclei, we pooled hippocampus aliquots from 5 female replicates and 5 male replicates within each age/genotype combination, resulting in 12 final pools (3 ages, two genotypes and two sexes). Tissue pools were homogenized separately in 2 mL chilled NIB (10 mM HEPES, pH 7.2, 10 mM NaCl, 3 mM MgCl_2_, 0.1% IGEPAL [v/v; Sigma-Aldrich, Cat#I8896], 0.1% Tween-20, and 1x protease inhibitor [Roche, Cat#11873580001]) and 10mM D(+)-Glucosamine hydrochloride [Sigma Aldrich G1414] in a 7 mL dounce-homogenizer on ice for 10 min. The homogenate was then strained through a 35 µm strainer and counted using a hemacytometer and trypan blue. The samples were centrifuged at 500 × g for 10 min at 4°C. Samples were aspirated, resuspended in ice-cold PBS-BSA buffer (0.5%. Bovine serum albumin (BSA), in 1x PBS, and 30mM D(+)-Glucosamine hydrochloride) to obtain 50,000 nuclei/5 ul of PBS-BSA-glucosamine. We tagmented 50,000 nuclei for each condition by adding 5ml ETB3 and 5 ml of an individual Tn5 (**Table 2**) at 37°C for 1 hour (f/c of D-glucosamine was 10mM) and iced 5 min. Nuclei were spun down (3 min, 500xg, 4°C) and washed in 1.5 mL TMG buffer (36% TAPS premix (4X TAPS-TD buffer (132 mM TAPS (N-[Tris(hydroxymethyl)methyl]-3-aminopropanesulfonic acid, Sigma-Aldrich, Cat#T0647-100G) pH=8.5, 264 mM potassium acetate, 40 mM magnesium acetate), 64% glycerol)/SCALE wash buffer twice. Nuclei were counted with a hemacytometer and trypan blue. Samples were multiplexed and processed as written in Step 2 (Gem Generation and Barcoding) of the 10x Genomics Single Cell ATAC v2 Kit protocol. 15 ml of each multiplexed pool of nuclei was added to 60 ml of the 10x Master Mix. We proceeded with the 10x protocol. For Step 4.1.c (Sample Index PCR), we substituted Sample Index N, Set A Reagent – with a ScaleBio S700 index primer compatible with the ScaleBio tagmentation (**Table 2**). Libraries were processed as described in the 10x Genomics Single Cell ATAC v2 Kit. They were quantified via the Qubit dsDNA High Sensitivity assay (Thermo Fisher Q32851) and via the Agilent Tapestation 4150 D500 tape (Agilent 5067–5592). Libraries were sequenced on the Illumina NextSeq2000 for 650 pM with a P2-200 flow cell (Illumina Inc., 20046812). ScaleBio tagmented libraries were sequenced paired-end with 85 cycles for read 1, 125 cycles for read 2, 8 cycles for index 1, and 16 cycles for index 2.

**Table 2.**
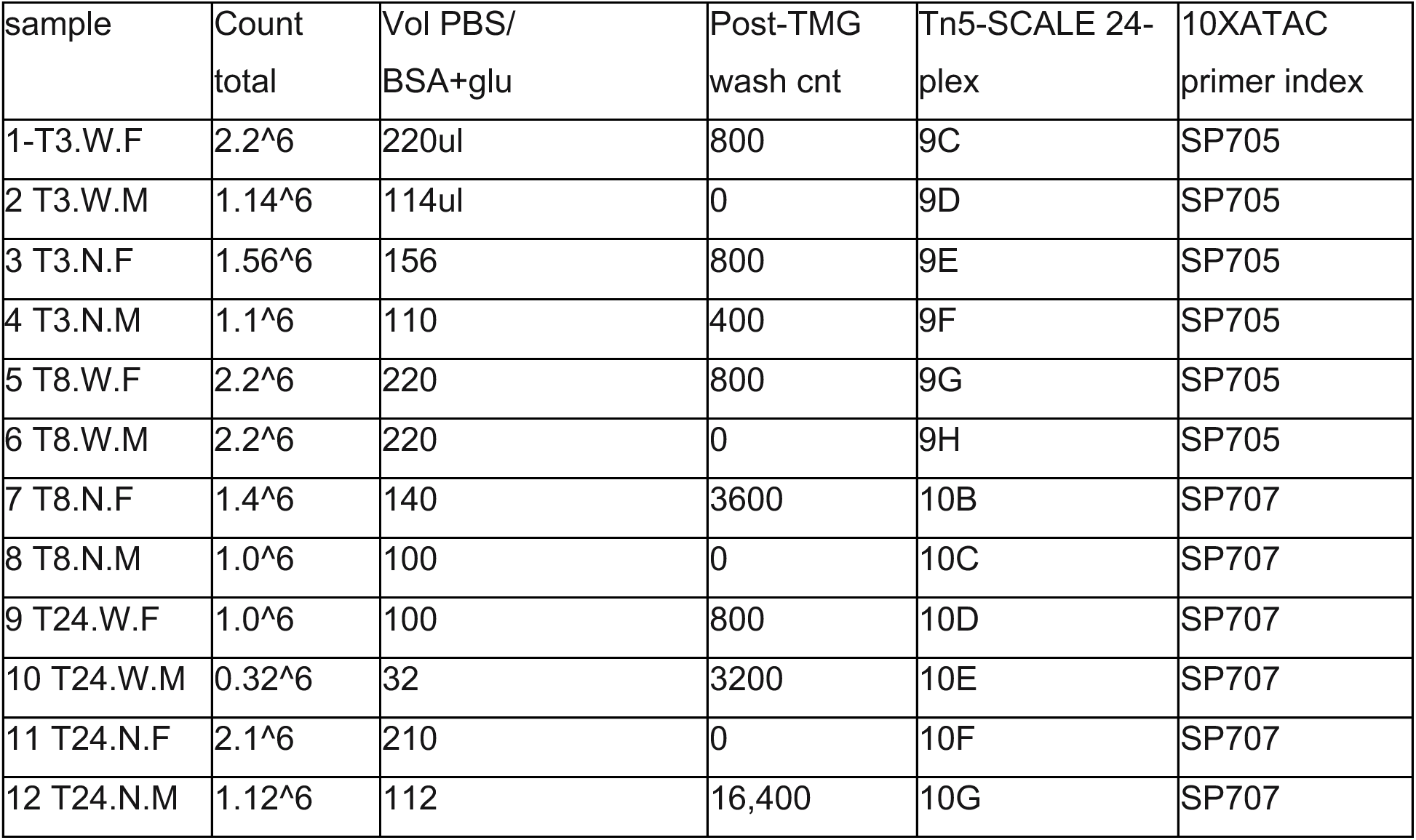
scATAC-seq library information. Primer index sequences were SP705:GGTCCAGACAAGGTCTACCTTGTGTTGAACAC, SP707:TCGGTACAGGAACGGTCATTGCTGATCCATAC

Raw scATAC-seq data was processed using the previously described scitools package (Sinnamon et al. 2019). Briefly, the sequencing reads were demultiplexed, and the read names were replaced with the cell barcode and a unique identifier. Reads were then mapped to the mouse genome (mm10) using BWA-MEM (Li and Durbin 2009). The resulting BAM files were filtered to remove low quality and duplicated reads. Additionally, we removed all barcodes with less than a specified number of reads. We used the ArchR (v 1.01) package (Granja et al. 2021) to filter cells with TSS enrichment < 2, unique fragments < 1000 and doublets. Using ArchR, iterative LSI dimension reduction and clustering was performed using the 500bp tile matrix, producing 21 clusters that were visualized via UMAP. Gene activity scores were determined and used to find marker genes for each cluster. Cell clusters were manually annotated via manual curation using top marker genes and publicly available data (Li et al. 2021; O’Connell et al. 2023). Out of the 21 clusters, four (clusters 8, 9, 10, and 19) could not be confidently classified and were broadly called “unclassified”. Clusters 9 and 19 were presumed to represent technical noise due to their UMAP distribution and lack of distinct marker genes (**Supplementary Table 1**). Clusters 8 and 10 were labeled as *Trpm3*-defined and *Grm4*-defined, respectively, based on their top marker genes (**Supplementary Table 1**). Cell composition of each sample pool was determined as percentage of total cells in the pool belonging to any cluster. Next, peaks were called at the pseudo-bulk level using all cells grouped by age/genotype combination, and cell-type. This was accomplished by using the addGroupCoverages function with parameters minCells = 30, maxCells = 100, minReplicates = 2, maxReplicates = 3, sampleRatio = 0.8 along with the addReproduciblePeakSet functions with parameters peaksPerCell = 500 and minCells = 20. Marker peaks/genes for each cell type and age/genotype combination were determined. archNext, we focused only on excitatory and inhibitory neurons (granule neurons were included as excitatory neurons). Peaks were called once again at the pseudo-bulk level grouped by age/genotype and cell type. Parameters for the addGroupCoverages function were minCells = 20, maxCells = 100, minReplicates = 4, maxReplicates = 8, sampleRatio = 0.8, and parameters for addReproduciblePeakSet were peaksPerCell = 500 and minCells = 20. Pairwise comparisons were performed using ArchR’s getMarkerFeatures to identify regions with significantly different chromatin accessibility between WT and *App^NL-G-F^*at each age, within excitatory neurons and inhibitory neurons.

### Bulk RNA-seq library preparation and analysis

Total RNA was extracted from 60 fresh frozen hippocampus aliquots using the NEB Monarch Total RNA Miniprep Kit (Cat T2010S, New England Biolabs). Briefly, samples were weighed and pulverized in a bead basher with DNA/RNA protection buffer. Pulverized samples were incubated with Proteinase K at 55 °C for 5 mins, followed by a 3 min centrifugation at 16,000 g. Supernatants were mixed with lysis buffer and centrifuged in gDNA removal columns. The flow through was mixed with 100% Ethanol and centrifuged in RNA columns at maximum speed. The columns were incubated with DNAse I/DNAse I buffer for 15 min at room temperature, followed by addition of priming buffer and centrifugation. Finally, the column was washed twice with a wash buffer and the RNA was eluted in water. RNA concentrations were measured with Nanodrop (Thermofisher) and integrity was confirmed (RIN > 7) with the Bioanalyzer RNA kit (Agilent). Samples were stored at −80 °C until further use. RNA-seq libraries were generated from 1ug of total RNA, beginning with an rRNA depletion of samples using the NEBNext rRNA Depletion Kit v2 for human/mouse/rat (New England Biolabs) according to manufacturer’s protocol. Samples were then purified with NEBNext RNA Sample Purification Beads (New England Biolabs). The RNA was then fragmented, primed, and reverse transcribed using the NEBNext Ultra II Directional RNA Library Prep Kit for Illumina (New England Biolabs). Libraries were prepared using the NEBNext Ultra II (New England Biolabs), with size selection to 300 bp using Ampure beads (Beckman Coulter). The final libraries were quantified using Qubit (Thermofisher) and single-end sequenced on the Illumina Nova-Seq 500 Platform at the Massively Parallel Sequencing Shared Resource (MPSSR) at Oregon Health and Science University (OHSU).

Raw reads were QC’d using FastQC (Andrews et al. 2019) and aligned to the mm10 genome using STAR (Dobin et al. 2013). The raw gene counts table was generated using custom bash scripts and filtered by the filterByExpr function from edgeR (Robinson et al. 2010) with default parameters and defining group membership as the combination of age and genotype. DESeq2 (Love et al. 2014) was used to perform differential analysis. Pairwise comparisons were performed to compare gene expression between wild-type and *App^NL-G-F^* at each age separately with sex as a covariate. The LRT test from DESeq2 was used to find genes with significant ageXgenotype interaction, followed by clustering according to the expression profile pattern using the degPatterns function from DEGreport (Pantano L 2024).

### Whole genome bisulfite sequencing (WGBS) library preparation and analysis

DNA from 60 hippocampus samples was extracted using the NEB Monarch Genomic DNA Purification Kit (Cat T3010, New England Biolabs) according to manufacturer’s protocol. Extracted gDNA was quantified with Qubit (Thermofisher) and Nanodrop (Thermofisher) and stored at −80 °C. DNA from 60 blood samples was extracted using the Purgene Genomic DNA Purification from Blood kit (Cat D-5500, Qiagen) based on a modified version of the Puregene DNA purification from blood protocol (Gentra Systems). Briefly, cell lysis solution was added directly to the frozen blood at a 5:1 ratio and incubated at 55 °C overnight, followed by 40 min or 1,000 rpm agitation. The samples were cooled on ice and protein precipitation solution was added (1:3 ratio) followed by 1,000 rpm agitation and 16,000 g centrifugation for 1 min. The lysate was moved to a 1:1 ratio of isopropyl alcohol and centrifuged to pellet the DNA. Pellets were washed with 80% ethanol, air dried, and resuspended at 65 °C with 1,000 rpm agitation. The gDNA samples were then quantified with Qubit (Thermofisher) and Nanodrop (Thermofisher) and stored at −80 °C. To generate whole genome bisulfite sequencing (WGBS) libraries for both hippocampus and blood,100 ng of gDNA was randomly sheared per sample with a Bioruptor Pico Sonicator (Diagenode) at 30:30 on/off for 15 cycles. Libraries were prepared with the NEBNext Ultra II Modules (New England Biolabs) and the NEBNext Methylated Adaptor (New England Biolabs), with size-selection to 200bp using Ampure beads (Beckman Coulter). Bisulfite conversion and subsequent cleanup was performed with the EZ DNA Methylation-Gold Kit (Zymo Research). The converted libraries were PCR amplified using NEBNext Q5U polymerase and the NEBNext Multiplex Oligos for Illumina (New England Biolabs) for unique library barcoding. The final libraries were quantified using Qubit (Thermofisher) and then normalized and multiplexed for single-end sequencing on the Illumina Nova-Seq 500 Platform.

The blood and the hippocampus WGBS datasets were analyzed separately, but similarly. Raw sequencing reads were trimmed with TrimGalore (Krueger), and then aligned to the mm10 reference genome with Bismark (Krueger and Andrews 2011) using default parameters. Deduplication was performed with deduplicate_bismark, followed by bismark_methylation_extractor with parameters – ignore 2, --ignore_r2 2, and --ignore_3prime_r2 2. Differential methylation analysis was performed with methylKit (Akalin et al. 2012) using the coverage files from Bismark methylation extractor data as input (overdispersion = “MN” and test = ”Chisq”). The genome was tiled into non-overlapping 1kb tiles, that were merged based on the following parameters: CpG coverage threshold ≥ 5, covered CpGs in the tile ≥ 10, and tile present in ≥ 3 samples per group. Sex was included as a covariate.

Overlap with gene features was determined using genomation (Akalin et al. 2015), with default settings. Intersection with the mouse cerebrum atlas of gene regulatory elements (Li et al. 2021) was done using Bedtools (Quinlan and Hall 2010). To investigate the distance between DMRs and DEGs, we used the closest-features program *(--closest --delim ’\t’ --dist*) from the BEDOPS toolkit (Neph et al. 2012) to calculate distance of each DEG from the closest DMR. These observed distances were then compared using the Wilcoxon signed-rank test against distances obtained against a randomly shuffled set of DMRs, generated by BEDtools shuffle (*-chrom -noOverlapping*) (Quinlan and Hall 2010).

### Gene Ontology term analysis

Gene ontology (GO) term analysis of differentially expressed genes (DEGs) from RNA-seq data was conducted using EnrichR (Chen et al. 2013; Kuleshov et al. 2016; Xie et al. 2021) with default settings. A subset of significant pathways (q < 0.01) was visualized across time points in dot plots using the ggplot2 library (Wickham 2016) in R. To assess and visualize the enrichment of biological functions associated with differential scATAC-seq peaks and WGBS DMRs, which include both coding and non-coding regions, we utilized gProfiler (Kolberg et al. 2020). The gProfler analysis were performed on the mm10 background, with an ordered query (sorted by adjusted p-values from most to least significant), using a 0.05 significance threshold based on the Benjamini-Hochberg false discovery rate correction. Dot plots of a subset of significant biological pathways were generated using ggplot2 (Wickham 2016) in R.

## DATA ACCESSIBILITY

Raw and processed RNA-seq, scATAC-seq and WGBS data generated as part of this study are available of the NCBI Gene Expression Omnibus (GEO; https://www.ncbi.nlm.nih.gov/geo/).

## AKNOWLEDGEMENTS

We thank Dr. Saido at the Riken Institute for sharing the *App^NL-G-F^*mouse model. Libraries were generated by the KCVI Epigenetics Consortium at OHSU and sequenced at the OHSU Massively Parallel Sequencing Shared Resource and at Novogene Corporation. Data analysis carried out in this study used computational infrastructure supported by the Office of Research Infrastructure Programs, Office of the Director, of the National Institutes of Health under Award Number S10OD034224. The content is solely the responsibility of the authors and does not necessarily represent the official views of the National Institutes of Health. This work was supported by the NIA award 1R21AG065681-01A1 awarded to LC and partially supported by R21 AG079158-01A1 and DOD HT94252410812 awarded to JR.

